# Deforestation in Belize-What, Where and Why

**DOI:** 10.1101/2020.01.23.915447

**Authors:** Hollie Folkard-Tapp

**Affiliations:** University of Southampton

**Keywords:** Forest Loss, Belize, Remote Sensing, Protected Areas, Deforestation Drivers, Vegetation Types

## Abstract

The tropical forests of Belize are a dynamic environment influenced by much disturbance throughout their natural history. In recent times, logging and agriculture have once more become the dominant drivers of land use change, yet the Belizean forests remain some of the most complete swathes of forest in the region. However, it is largely unknown how the abundance of specific forest types has changed. This study aims to critically assess whether valuable habitat types have been disproportionately affected by deforestation between 1986 and 2018. ENVI Classic 5.5 was used to calculate the NDVI values of Landsat imagery, with values over 0.6 deemed to indicate primary forest. ArcMap 10.6 was used for detecting change in forest cover between 1986 and 2018, in addition to digitising colonial maps. Results show variation in the percentage of forest lost between habitat types, with a bias towards forests fragmented by or replaced by agriculture, particularly in the north and along the Guatemalan border. The protected area network was found overall to have little influence on deforestation, though this varies between reserves. Of the three forest types experiencing the highest percentage of forest loss, only Ensino-Pixoy occurs exclusively outside protected areas. Cohune Santa-Maria and Sapote-Mahogany are found at least partially within protected areas and experience loss within them. Analysis of IUCN ratings of forest species revealed data deficiency, particularly concerning taxa indigenous to Mesoamerica, against which this study recommends further vegetation studies. to allow recommendations for protection of particularly threatened forest types. A review of management strategies in protected areas, especially the Caracol Archaeological Reserve, is necessary to avert the worrying deforestation trends this study has identified. The country-wide deforestation data can be used to advise future forest policy as well as calculation of national carbon stocks or determining integrity of ecosystem services.

## Introduction

Tropical forests are intrinsic to many global ecosystem services (Wunder 2005; Young, 2008). Deforestation does not only threaten their ability to provide cultural, provisioning and regulating services, but also undermines the supporting services needed to maintain all others (Patterson, 2016; Tadesse *et al*., 2014). Changes to forest composition and structure are the primary cause of biodiversity loss worldwide (Sanchez-Azofeifa *et al*., 2001), and cause destabilisation of climate system dynamics and atmospheric composition (Daily, 1997). Globally, tropical deforestation is responsible for approximately 20 per cent of global greenhouse gas emissions (IPCC, 2013). With increasing global commitments to mitigate climate change, monitoring of forest cover is gaining importance due to its direct relevance to carbon stocks (Rosenqvist *et al*., 2003). There is also a need to understand local drivers of deforestation in the tropics to ensure effective policy interventions at this scale (Patterson, 2016).

To develop reliable baselines for the forestry sector, which can then be used to combat deforestation and associated impacts, three fundamental questions (Kalacska *et al*., 2008) need answering: *“What is the initial extent of the forest? What type of forest is there? What is the rate of change in forest extent?”*

Due to the lack of standard methodologies capable of providing accurate data for different tropical forest types, establishing these baselines is difficult (Foster *et al*., 2002; Subak, 2000). The tropical forests of Belize are a dynamic environment that has been influenced by various levels of disturbance throughout their natural history (Bridgewater, 2012), and therefore have a very varied architecture. In the 20^th^ century, logging and agriculture became the dominant drivers of land use change and forest loss, along with cattle ranching and urban growth (Horwich & Lyon, 1990). After British colonisation in 1862, forestry flourished, supporting the economy until British Honduras attained self-governance in 1964 (Bridgewater, 2012; Bolland, 1992). After this point, and the granting of full independence in 1981, population increase was associated with a rise in agriculture, both through traditional slash-and-burn *Milpa* methods and modern intensive farming (Bridgewater, 2012; Patterson, 2016).

Shortly before this shift to agriculture, in 1959, the British Honduras Land Use Survey Team of the Colonial Office published an assessment of vegetation types to provide advice for agricultural development in the Colony (Wright *et al.,* 1959). This research provides a highly detailed record of habitat architecture pre-intensive farming, and so acts as a baseline of semi-natural forest extent.

Wright *et al*. (1959) prepared these maps using aerial photographs and ground survey data from multiple sources including the Director of Land Surveys in British Honduras and the British Honduras Gulf Oil Company. At the time, the work was considered necessary for two reasons; to estimate the local climate in areas where records had not been kept, and to indicate soil fertility in areas with no recorded farming experience. The resulting map that therefore ‘portrays the original, untouched pattern of the vegetation that soils could be expected to support’ (Wright *et al*., 1959). Where ecological reality deviated from the predictions, the influence of fire or ‘very old, unrecorded agriculture’ was cited (Wright *et al*., 1959). This includes Mayan agricultural influence, and therefore the baseline is set not before human manipulation of the ecosystem, but before the advent of modern intensive farming practices.

Despite these colonial efforts to maximise the farming potential of Belize, the region retains some of the most complete swathes of forest in Central America (Bridgewater, 2012). Latest official figures state that Belize retains 79% forest cover (FAO, 2004), though this includes ‘other wooded areas’ which do not fit this study’s definition of forest. More recent studies estimate it to be closer to 60% (Meerman *et al*., 2010; Cherrington and Ek, 2010, Voight *et al*., 2019). This is a testament to the ecological resilience of the area, despite – or perhaps because of – the history of disturbance through the ages. Despite this resilience, the forest is showing signs of both human and natural pressures from recent years. Latest, but outdated, official figures state annual rate of deforestation to be 36000 ha pa (FAO, 2004); twice the Central American average - 2.3% vs 1.2% respectively (Young, 2008). At this rate, forest cover was predicted to decrease to 58% by 2020 and be completely lost by 2050 (Young, 2008).

Differences between the outdated FAO forest cover estimations and those calculated by more recent remote sensing projects show the importance of further research and validation via these methods, further highlighting the need to establish reliable baselines and answering the remainder of Kalacska’s (2008) fundamental questions. In addition to the uncertainty surrounding the exact rate of forest loss, it is unknown how the distribution of specific forest types has been affected. There is a significant gap in the literature covering this phenomenon, with few attempts to differentiate between primary and secondary forest and even less investigating species composition on a country-wide scale.

This paper aims to determine baseline abundance for specific habitat types pre- and post-intensive agriculture and calculate forest loss, critically assessing whether the distribution and area of ecologically valuable habitats have been influenced by deforestation in the last 30 years. In order to achieve this, Wright *et al*. 1959 maps have been digitised to allow calculation of the pre-intensive agricultural extent of each forest type defined in that study. The earliest post-agricultural intensification Landsat imagery is used to determine a more accurate baseline of forest cover in mainland Belize, against which latest data can be compared to calculate the rate of deforestation affecting different habitat types, within and outside of protected areas. Finally, IUCN Red List classifications are used to investigate how habitats supporting high-value species have been affected by deforestation.

## Methods

As in previous studies of vegetation cover in Belize (Meerman *et al*., 2010; Ross, 2011), this project defines forest as “closed canopy, mature natural broadleaf forest”. This definition was chosen primarily due to its ability to be classified through remote sensing imagery. NDVI in particular is capable of determining areas of this forest classification. Tropical forests appear as regions with reflectance values above 0.6 (Wittich and Hansing, 1995; Weier and Herring, 2000). ENVI Classic 5.5 is a program with the capability to process Landsat imagery to calculate NDVI automatically. This study utilises that functionality, as well as the change detection capabilities of ESRI’s ArcMap 10.6.

This methodology follows three main steps: Calculation of forest cover in 1986 and 2018 using satellite imagery; Estimation of forest cover in 1959 through analysis of historical maps; and change detection between the timeframes.

### Data collection and pre-processing

Atmospherically corrected Landsat data was downloaded from the USGS EarthExplorer website (https://earthexplorer.usgs.gov/) in November 2018. The study aimed to use images from two dates as far apart as possible, and on the same day of the year, while not being unreasonably compromised by cloud cover. The dates that suited these parameters best are 07/04/1986 and 07/04/2018. Three spatial data frames were required for each year to cover the entirety of mainland Belize-Rows 47, 48 and 49 of Path 19, which correspond to northern, central and southern Belize respectively. The 1984 images were collected by the Thematic Mapper (TM) aboard Landsat 5, the 2018 images by the Enhanced Thematic Mapper Plus (ETM+) aboard Landsat 7. Both these images are at a resolution of 30m and have comparable wavelengths.

Image pre-processing in ENVI consisted of refining, for each image, the cloud and water masks applied by USGS. All analysis was conducted at a spatial resolution of 30m, which can detect forest presence or absence down to less than one hectare, using only pixels which were free of clouds across both dates. Full details of image pre-processing in ENVI, as well as subsequent analysis and digitising of the historical maps in ArcMap, are shown as an IDEF0 process flow (Appendix 1). After applying these corrections, NDVI was calculated using the equation:

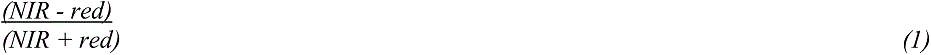

where *NIR* is Landsat Band 4 and *red* is Landsat Band 3.

The resulting NDVI image was imported to ESRI ArcMap 10.6, where images were re-projected to UTM NAD 1927 Zone 16N; as used by the scanned paper map.

### Post-processing

Polygon feature classes representing forest and non-forested areas for each year were created for ease of comparison with the digitised vegetation types proposed by Wright *et al*., 1959 (Figure 1). Areas with an NDVI value of >0.6 were deemed to fit this study’s definition of ‘forest’ which describes broadleaf vegetation exclusively (Wittich and Hansing, 1995).

**Figure 1.**
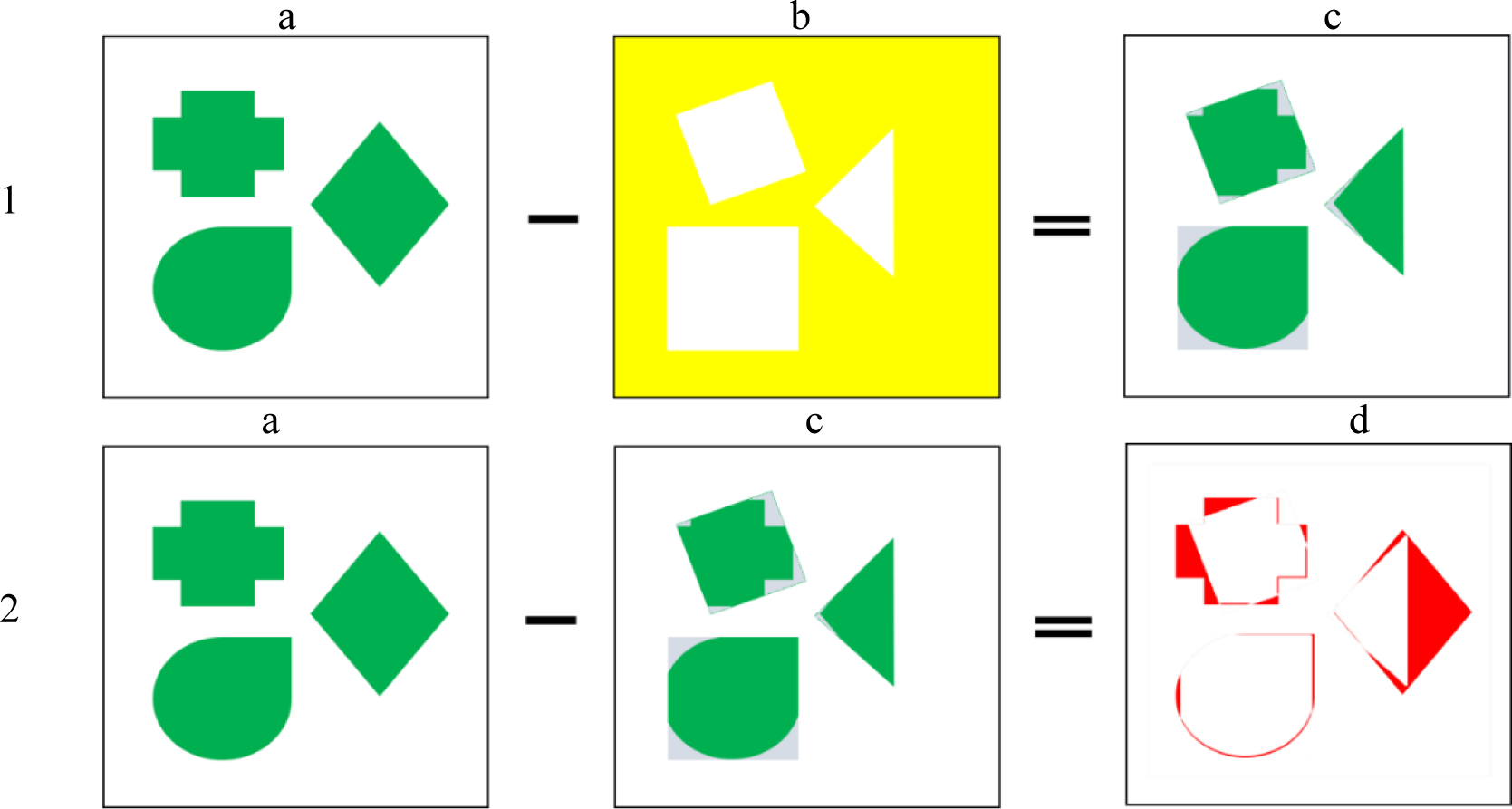
Schematic of original forest remaining and forest loss calculation. a) 1986 forest (green), b) 2018 non-forest (yellow), c) remaining forest (green) and secondary growth (grey), d) forest lost (red). *Image analysis (change detection)*

The 2017 Belize Ecosystems Map (Meerman and Clabaugh, 2017), based upon UNESCO system for Central American ecosystems, was used to assess the accuracy of both the 1959 vegetation map and the NDVI forest classification. The 11 broad categories proposed in 1959 largely correlated to the UNESCO forest types, which in turn are reflected in the NDVI-generated broadleaf forest extent.

To calculate the regions of 1986 or ‘original’ forest that remained in 2018 (Figure 1.1), the 2018 non-forest polygons were erased from the 1986 forest polygons. To show the regions that had been lost, this ‘remaining’ output feature class was removed from the 1986 forest (Figure 1.2).

This methodology ensures that vegetation regrowth with an NDVI value of >0.6 that has occurred since 1986 is not included in the feature class of current forest cover, as it is neither mature or natural, and therefore does not meet this study’s definition of ‘forest’.

To determine deforestation levels between 1986 and 2018 within each vegetation type, the countrywide forest/non-forest polygons were clipped to create feature classes comprising polygons representing each 1958 habitat fragment. The sum of these areas resulted in a value for the number of hectares of each forest type in 1986 and 2018. The difference between these two provides a clear record of decline and fragmentation of Belize forest stocks.

## Results

Between 1986 and 2018, Belize’s forest stocks declined by 28.4%, with 235,432 ha experiencing deforestation and representing a land cover change of 4% across the country. Some habitat types were disproportionately affected, with Sapote-Mahogany suffering the greatest absolute decline and Ensino-Pixo experiencing the highest percentage loss. Protected areas were found to experience similar levels of deforestation to those without designations. 35% of forest loss occurred in this protected 38% of land area. Chiquebul-Cherry, found almost exclusively within protected areas, remains the most abundant forest type across all timeframes.

Digitising the 1959 map (Figure 2) resulted in 2,125,074 ha of land in mainland Belize classified as one of 76 vegetation types across 11 broad categories (Appendix 2). Due to masking of pixels affected by cloud cover and striping caused by faults in the ETM+ sensor’s Scan Line Corrector (SLC) (Chen *et al*., 2011), the total area sampled from the Landsat data was 1,371,662ha, 65% of the entire data frame. For this reason, and because the original maps did not consider existing settlement or roads, direct comparisons between forest area in 1959 and modern records are not appropriate. Therefore, only changes occurring between 1986 and 2018 can be reliably quantified. These results can be visualised in two manners (Meerman *et al*., 2010).

1. Strictly as forest cover for each year (Figure 3)
2. As areas where change was detected (Figures 3 and 4)

**Figure 2.**
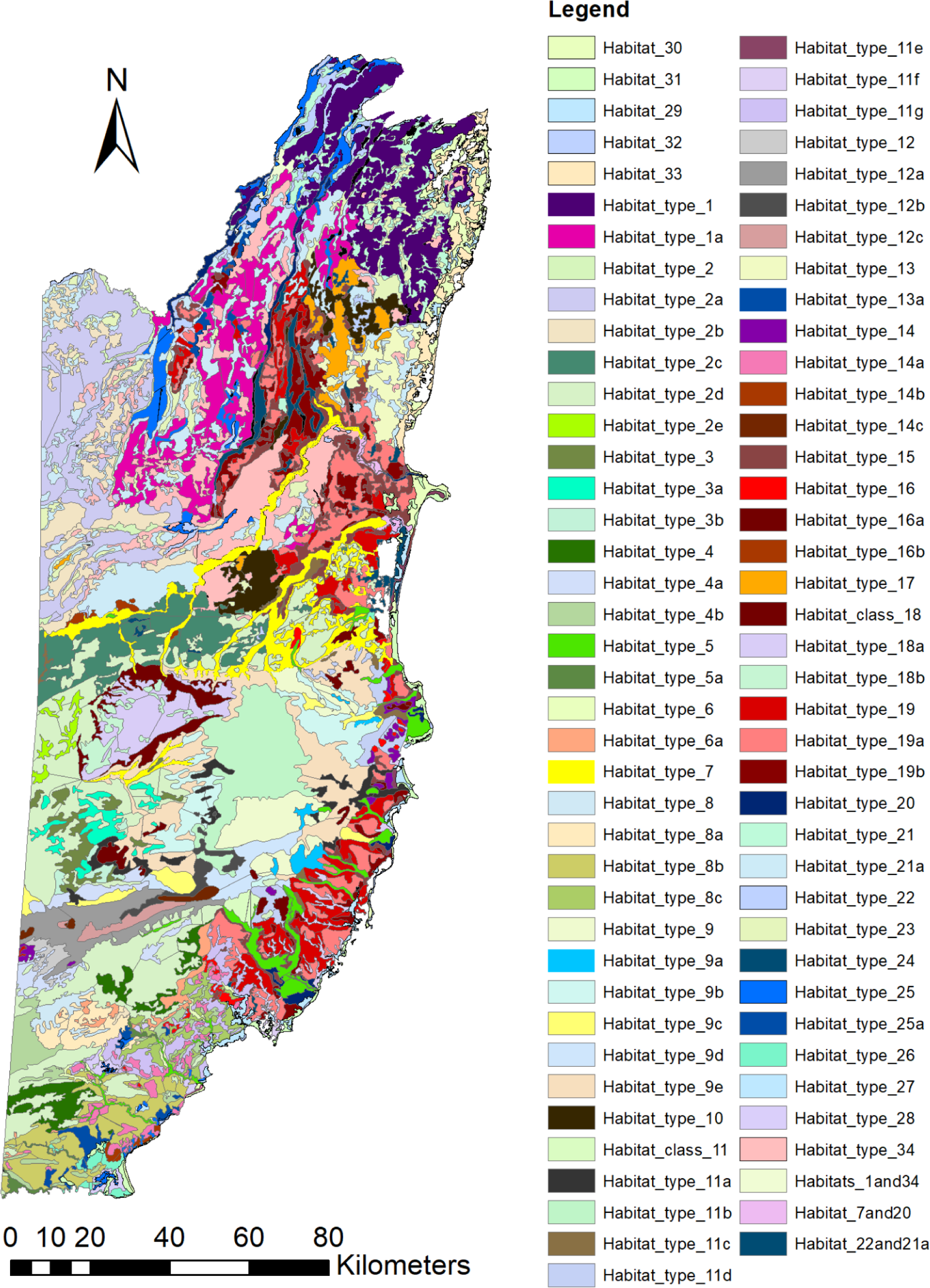
Digitisation of Wright et al., (1950) Natural Vegetation Map (Sheets 1 and 2), <u>completed in ArcMap10.6. Habitats 30-33 are found within the littoral zone.</u>

**Figure 3.**
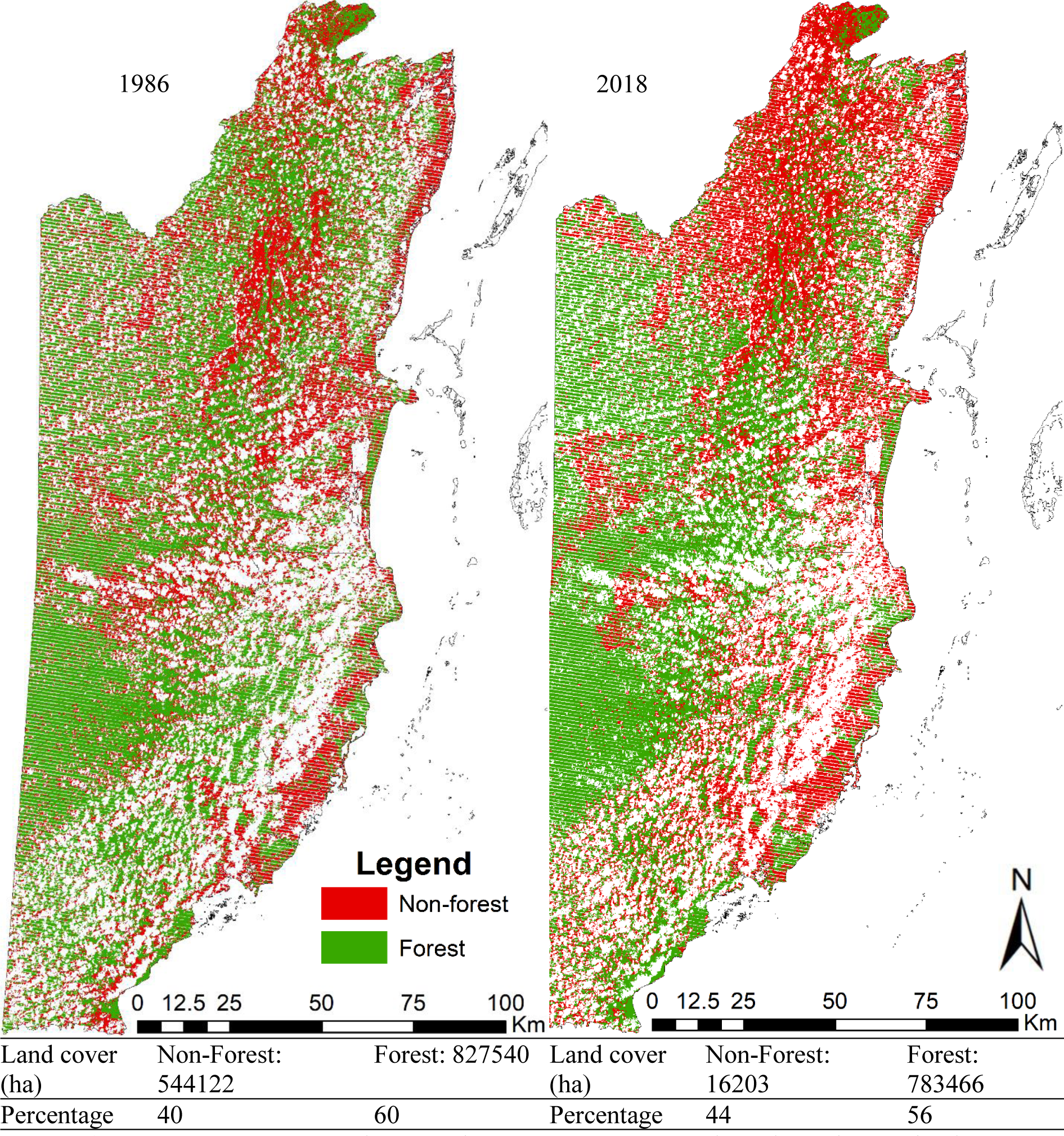
Forest cover in 1986 and 2018. White areas represent no data, due either to cloud cover or the striping seen in 2018 Landsat ETM+ data.

The first method of visualisation is useful as an overview of vegetation distributions but does not account for areas of secondary growth that have occurred since the first study date.

From 1986 to 2018, 219,129 ha of secondary forest growth was observed, mainly outside of the heavily agricultural northern areas. This study focusses on visualising forest loss as areas of recorded change to detect and exclude these immature stands from calculations of mature forest stocks. From 1986 to 2018, a total of 235,432 ha of forest were lost, representing a decline of 28.4% across all vegetation types, or 0.89% annually (Figure 4).

**Figure 4.**
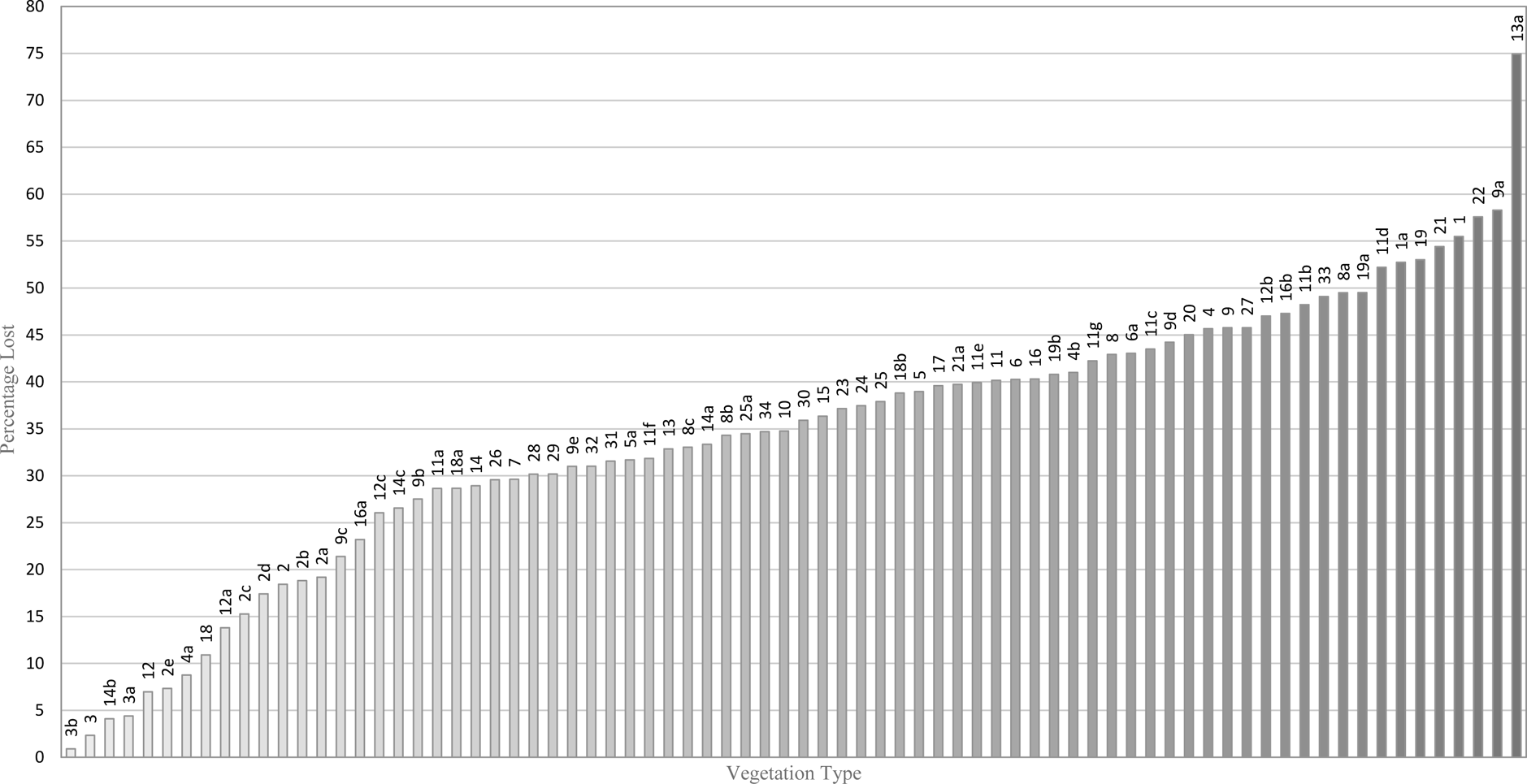
Percentage of forest lost by vegetation type between 1986 and 2018.

There is great variation between vegetation types, with less than 5% of Semi-evergreen, 80-100ft high on limestone forests being lost (Types 3, 3a and 3b), whereas others have lost over 50%, such as Sapote-Mahogany (Type 1), Cohune Santa-Maria (Type 9a) and Pucte-Chechem (Type 22). The area covered by Ensino-Pixoy forest (Type 13a), a form of transitional low broadleaf forest locally known as broken ridge, has declined by 1,615 ha, almost 75%, since 1986. This value falls high above the trend for deforestation percentages, especially when compared to the values for the other Broken Ridge assemblages (Habitat Types 13-15).

Between the eight habitat types experiencing over 50% loss (Table 1), there is pronounced variation between the species present and the size of the initial forest area. The two habitat types experiencing the same percentage loss, Cohune Santa-Maria and Pucte-Chechem, vary in absolute size by a factor of 10 in terms of absolute size.

**Table 1.** Habitat types experiencing the most deforestation as a **percentage** of their original size.

| Code | Main Forest Species | Other Notable Species | Size 1987 (ha) | Remaining 2018 (ha) | Total Loss (ha) | % Loss |
| --- | --- | --- | --- | --- | --- | --- |
| 13a | Ensino-Pixoy | Grape, Cojeton, Tastab, Botan Palm, Black Maya, Cutting Grass | 2155 | 540 | 1615 | 75 |
| 9a | Cohune-Santa Maria | Fig, Negrito, Star-Apple, Quamwood, Warrie-Cohune, Cojeton, *Heliconia* (Mahogany stumps) | 2072 | 864 | 1208 | 58 |
| 22 | Pucte-Chechem | White Wattle, Knock-Me-Back, Logwood, Redwood, Palmetto, Basket Tie-Tie | 10970 | 4653 | 6317 | 58 |
| 1 | Sapote-Mahogany | Negrito, Cabbage Bark, Turtlebone, Gombolimbo, Grande Betty, Fiddlewood, Grape, Glassywood, | 35223 | 15677 | 19546 | 55 |
| 21 | Pucte-Chechem | Sapote, Cockspur, Cibish, Salam, Gombolimbo, Madre de Cacao, Chucum | 10741 | 4896 | 5845 | 54 |
| 19 | Grass | Pine, Crabboe, Yaha, Oak | 5273 | 2477 | 2796 | 53 |
| 1a | Sapote-Mahogany | Pixoy, Cockspur, Cabbage Bark, Turtlebone, Give-and-Take, Thatch Palm | 35959 | 16993 | 18966 | 53 |
| 11d | Nargusta-Can't-help-it-Mahogany | Cohune Palm, Black Maya | 3436 | 1642 | 1794 | 52 |

Within this group, the two forest types that have experienced the highest loss in absolute terms are comprised mainly of Sapote-Mahogany assemblages. They also had the greatest initial size. 1a and 1, Sapote-Mahogany forests, were the 5^th^ and 6^th^ most abundant types in 1987, and remain 5^th^ and 7^th^ largest in 2018 respectively. Of the 482,985 ha of forest remaining in 2018, Chiquebul-Cherry, Botan-Chechem and Sapote-Silion remain the most abundant types (Table 2). Chiquebul-Cherry lost only 17% of its area, while Botan-Chechem lost almost 40%.

**Table 2.** Three most abundant habitat types by **absolute area** remaining in 2018

| Code | Main Forest Species | Other Notable Species | Size 1987 (ha) | Remaining 2018 (ha) | Forest Loss (ha) | % lost |
| --- | --- | --- | --- | --- | --- | --- |
| 2d | Chiquebul-Cherry | Ramon, Bullhoof, Sapote, Copal, Cedar, Fiddlewood, Mahogany | 82423 | 68069 | 14354 | 17.4 |
| 21a | Botan-Chechem | Pucte, Basket Tie-tie, Sapodilla, Grape | 50836 | 30635 | 20201 | 39.7 |
| 2a | Sapote-Silion | Mahogany, Spice, Mylady, Ramon, Give-and-take palm | 37098 | 29978 | 7120 | 19.2 |

To determine whether the initial size of the forest is a significant factor in determining forest loss, Spearman’s Rank tests found that initial forest size has a strong correlation with *absolute* forest loss (rs = .858, p = .001), but is not significantly correlated with the *percentage* of initial forest loss. The correlation between percentage loss and absolute loss in hectares is only moderately significant (rs = .347, p = .002). Comparing deforestation between habitat types is therefore most appropriate using percentage loss over absolute.

Chiquebul-Cherry forest (Type 2d), remains the most abundant across all timeframes studied. This low level of absolute forest loss is echoed in the percentage loss of only 17%. It occurs mainly in the Chiquebul Forest Reserve. Protected areas cover 36% of mainland Belize (Young, 2008). Of the deforestation that occurred during the study period, 833,61 ha was lost from these areas, representing 35% of all forest loss (Figure 6). This shows that the rate of deforestation in these protected areas is only slightly below the national average.

Wright *et al*. (1959) recorded 50 species as comprising the habitat types suffering over 50% loss. Of these, four are given conservation status by the IUCN. Both species of Fiddlewood are endangered, and Heliconia and Mahogany are Vulnerable (Ulloa Ulloa and Pitman, 2004). In addition to these, 20 other taxa have also been classified by the IUCN on Red List of Threatened Species, 6 of which require some form of conservation effort. (Table 3).

**Table 3.** All available IUCN Redlist classification of species found in Belize, as recorded by Wright et al., 1959.

| Local name | Latin name | IUCN rating | Uses |
| --- | --- | --- | --- |
| Fiddlewood | *Vitex guameri* | EN | General furniture |
| Fiddlewood | *Vitex kuylenii* | EN | General furniture |
| Calabash Tree | *Crescentia cujete* | VU | Edible and medicinal |
| Spanish Cedar | *Cedrela mexicana* | VU | Wood for cigar boxes |
| Heliconia | *Heliconia fredberryana* | VU | Ornamental |
| Mahogany | *Swietenia macrophylla* | VU | Medicine, agroforestry, Timber |
| Mountain palmetto | *Schippia concolour* | VU | Ornamental |
| Mylady | *Aspidosperma megalocarpon* | NT | Medicine (malaria), heavy carpentry, fuel |
| Wild tamarind | *Acacia dolichostachya* | NT | Medicine |
| Alsophila fern | *Alsophila myosuroides* | LC | Medicine (stops bleeding) |
| Bastard mahogany | *Carapa guianensis* | LC | Timber, medicine, insect repellent |
| Black Mangrove | *Avicennia nitida* | LC | Medicine |
| Cotton Tree | *Ceiba pentandra* | LC | Food, medicine, agroforestry, timber |
| Cypress | *Podocarpus guatemalensis* | LC | Timber, medicine |
| Fig | *Ficus glabrata* | LC | Soft timber, latex |
| Fig | *Ficus radula* | LC | Soft timber, latex |
| Logwood | *Haematoxylum campechianum* | LC | Medicine (anti-inflammatory), dye, essential oil |
| Monkey Apple | *Licania platypus* | LC | Fruit, timber (uncommon) |
| Ramon | *Brosimum alicastrum* | LC | Food, medicine, cheap timber, latex |
| Santa Maria | *Calophyllum brasiliense* | LC | Medicine, lamp oil, valuable timber |
| Barba Jolote | *Pithecolobium arboreum* | LC | Food, medicine, agroforestry |
| Dumb Cane | *Dieffenbachia sp.* | DD | Food, medicine, poison, ornamental |
| Waika Chewstick | *Symphonia globulifera* | DD | Fruit, medicine, resin, latex, dye, low-quality durable timber |

Across the seven forest types experiencing the least deforestation by percentage of 1986 area, below 10%, only seven taxa have been evaluated by the IUCN Red List. Both species of Fiddlewood are Endangered, Mahogany and Cedar are Vulnerable, Mylady is Near Threatened, and Ramon and Santa Maria are Least Concern. Of these seven forest types experiencing less than 10% loss, five are found almost exclusively within the Chiquebul forest reserve. No Chiquebul forest types experience over 50% loss, the highest being Chiquebul-Santa Maria forest types 4 at 47%, and 4b at 41% loss. In the fully-protected Chiquebul-Cherry (Type 2d) forest, the habitat covering the most significant area across all timeframes, three Red List species are present; Mahogany, Cedar and Fiddlewood.

## Discussion

The first stage of determining rate and cause of deforestation is establishing reliable baselines against which change can be measured (Kalacska, 2008). This study used Wright *et al*.’s 1959 Natural Vegetation maps to calculate a pre-industrial agricultural baseline of forest composition and coverage. Digitising the maps revealed that 2,125,074 ha of mainland Belize were classified by the British Honduras Land Use Survey Team in 1959, comprised of 76 vegetation types across 11 broad categories. These maps portray the original, untouched pattern of the vegetation, based mainly on what the local soils could be expected to support (Wright *et al*., 1959). They do not account for recent agricultural activity, settlements, forest fires or any other form of disturbance, whether human or natural. Many rivers and creeks are also missing from the map. For these reasons, it is apparent that the estimate of forest cover provided by the 1959 maps is an overestimation, and that the baseline it provides is not entirely reliable. In addition to this, the study area from 1986-2018 was limited by masking pixels affected by cloud cover and ETM+ striping. The total area sampled from the Landsat data was 1,371,662 ha, 65% of the area mapped by Wright *et al*. (1959). Direct comparisons between forest area in 1959 and Landsat-derived modern calculations are therefore not appropriate. Only changes in forest cover between 1986 and 2018 can be reliably quantified. The 1959 maps were therefore used mainly for the classification of the vegetation and their general distribution, answering the second of Kalacska’s fundamental questions for managing deforestation, ‘*what type of forest is there?’* (Kalacska, 2008). Although this register of forest species is not exhaustive, the information that exists is likely to be accurate, as Wright *et al*. used a method (Holdridge, 1950) of vegetation classification which they were confident of being suitable for both their level of skill and the time they had available for the study (Wright *et al*., 1959).

To answer the first of Kalacska’s questions (2008), *“What is the initial extent of the forest?”,* the earliest suitable Landsat data, April 1986, therefore serves as a post-agricultural intensification baseline. For comparison, the most recent images with low cloud cover from an anniversary date were chosen to indicate current forest cover in Belize. Calculating change between these two timeframes fulfils the final of Kalacska’s (2008) questions, concerning the “*change in forest extent”.* The loss of 235,432 ha of forest from 1986 to 2018 represents a decline of 28.5%, averaging 0.89% annually. This falls slightly above the Cherrington *et al*. (2010) calculations of 0.6% annual rate of forest loss between 1980 and 2010, with a total loss of 17.4% over the 30 years (Cherrington and Ek, 2010). Assuming both calculations are correct, the annual deforestation rate appears to have increased since 2010. Further studies calculating forest cover at interim dates would confirm this. Alternatively, the studies differ slightly in the methodology for distinguishing secondary forest growth. Cherrington’s (2010) method of using supervised classification to detect spectral differences between primary and secondary forest does not account for stands of fast-growing timber planted on previously clear-felled land that reached maturity within the span of the study period. This study does not consider this to be primary forest as its growth is due to human interference and will not provide the same ecosystem services as mature, natural forests (Poorter *et al*., 2016). Additionally, the species composition of these altered areas differs to that recorded by Wright *et al*. in 1959, and so cannot be compared against this data.

Due to its real-world management implications, absolute forest loss remains a relevant consideration when investigating reasons for the deterioration of certain forest fragments, particularly in small, isolated areas (Wyman, 2008). However, percentage loss was deemed the most appropriate measure for comparing of loss of each habitat type rather than absolute loss for ease of comparison (Emch 2008; Weishampel *et al*., 2012) and because it is not so heavily influenced by original fragment size (Figure 5). Fragmentation and the theory of island biogeography is well established as a cause of habitat loss (Harris, 1984; Meave, 1991; Redford, 1992; Wyman, 2008), and limiting the influence of this cause of deforestation in Belize allows other factors, such as species composition and incidence of protected areas, to be considered.

**Figure 5.**
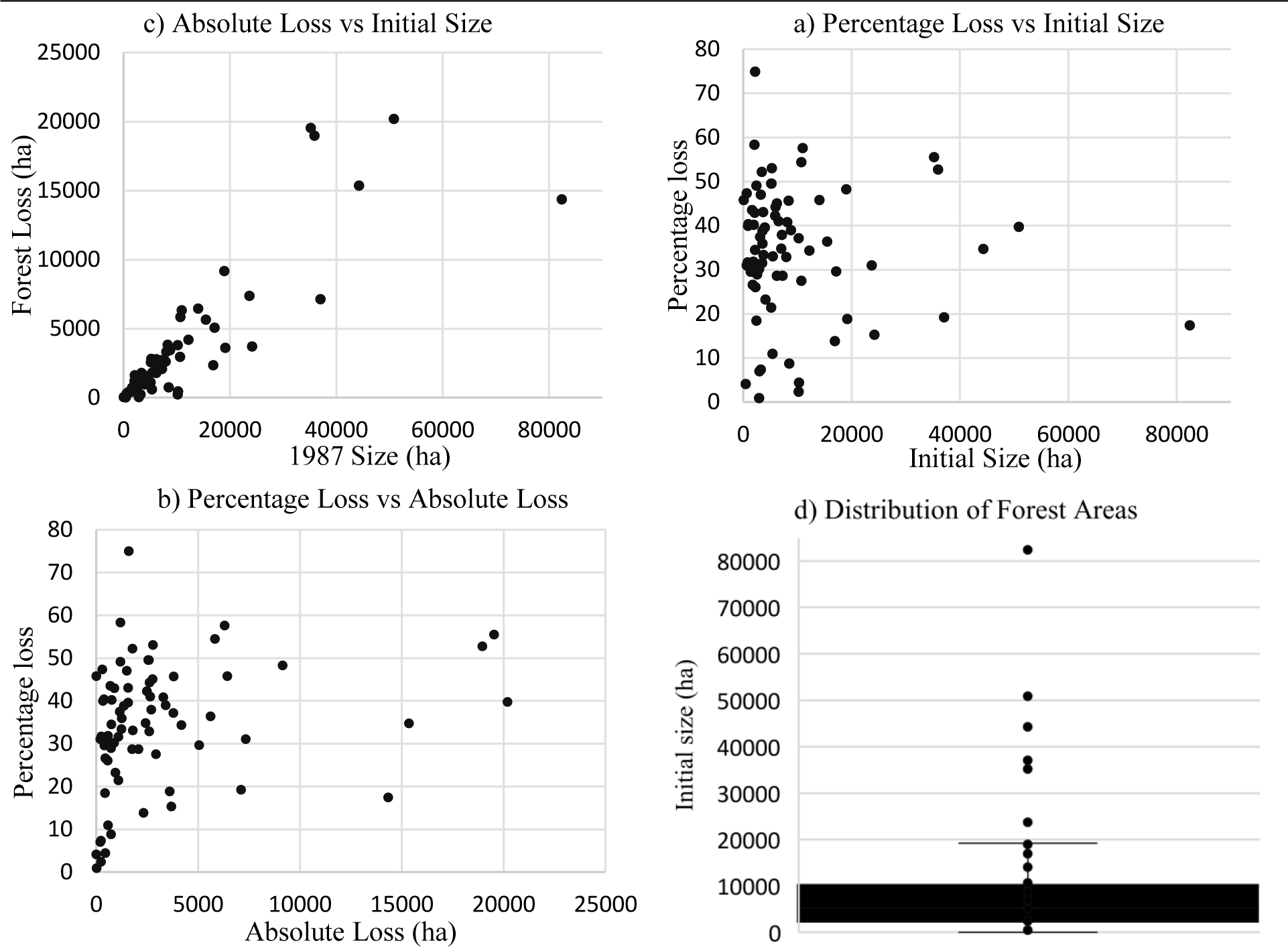
a) Correlation between percentage loss of forest against initial size of each habitat type. b) Correlation between the percentage of original forest loss and the absolute number of acres of each habitat type deforested. c) Acres of forest loss correlated against the initial size of the habitat type. d) Distribution of habitat types by initial size showing greatest variance in the upper quartile.

Although 38% of land is under some form of legal protection, only 13% is classified as biodiversity reserves; the rest being extractive reserves allowing the removal of timber, flora and fauna (Gonzales, 2007). Most protected areas are found towards the south of Belize as most of the north was already privately owned by the time the protected areas were allocated, first by the Forestry Trust in 1923, and then the Forest Department in 1936 (King *et al*., 1992). Much of the forest loss detected within these areas occurred on the western edge of the central protected zone, as well as in the northern Freshwater Creek Forest Reserve.

Freshwater Creek was created as a Crown Reserve in 1930, and received its current conservation status in 1960. It was found to be comprised mainly of Sapote-Mahogany and Sapote-Silion forest types (1 and 2) (Wright et al, 1959). The area was historically logged for Mahogany and Logwood, leading to a lack of commercial-sized timber remaining in the Eastern half by 1990, and no management plan was implemented to prevent further degradation (King *et al*., 1992).

Nationally, the highest levels of deforestation occur in the northern districts of Corozal and Orange Walk; which are characterised by extensive farming (Cherrington and Ek, 2010). The complex road network surrounding agricultural plots where once was Ensino-Pixoy forest (Type 13a) illustrates the expansion of the industry from 1959 to 2018 (Figure 7). Freshwater Creek Forest Reserve straddles both these districts. From 1980 to 2010, the area cleared for agriculture expanded by 65%, with sugarcane featuring strongly as a cash crop (King *et al*., 1992). Mennonite communities form a large labour force which also produces rice, beans, corn and cattle, leading to the highest rates of land clearance, land degradation, and unregulated use of pesticides in the country (Ministry of Forestry, Fisheries and Sustainable Development, 2014). The deforestation detected in this region between 1986 and 2018, such as the loss of Sapote-Mahogany habitat (Type 1) inside and outside of protected areas (Figure 7), is therefore likely to be a combination of remaining high-value timber removal and rapidly expanding agriculture driven by population increase (Emch, 2008; Patterson, 2016).

**Figure 6.**
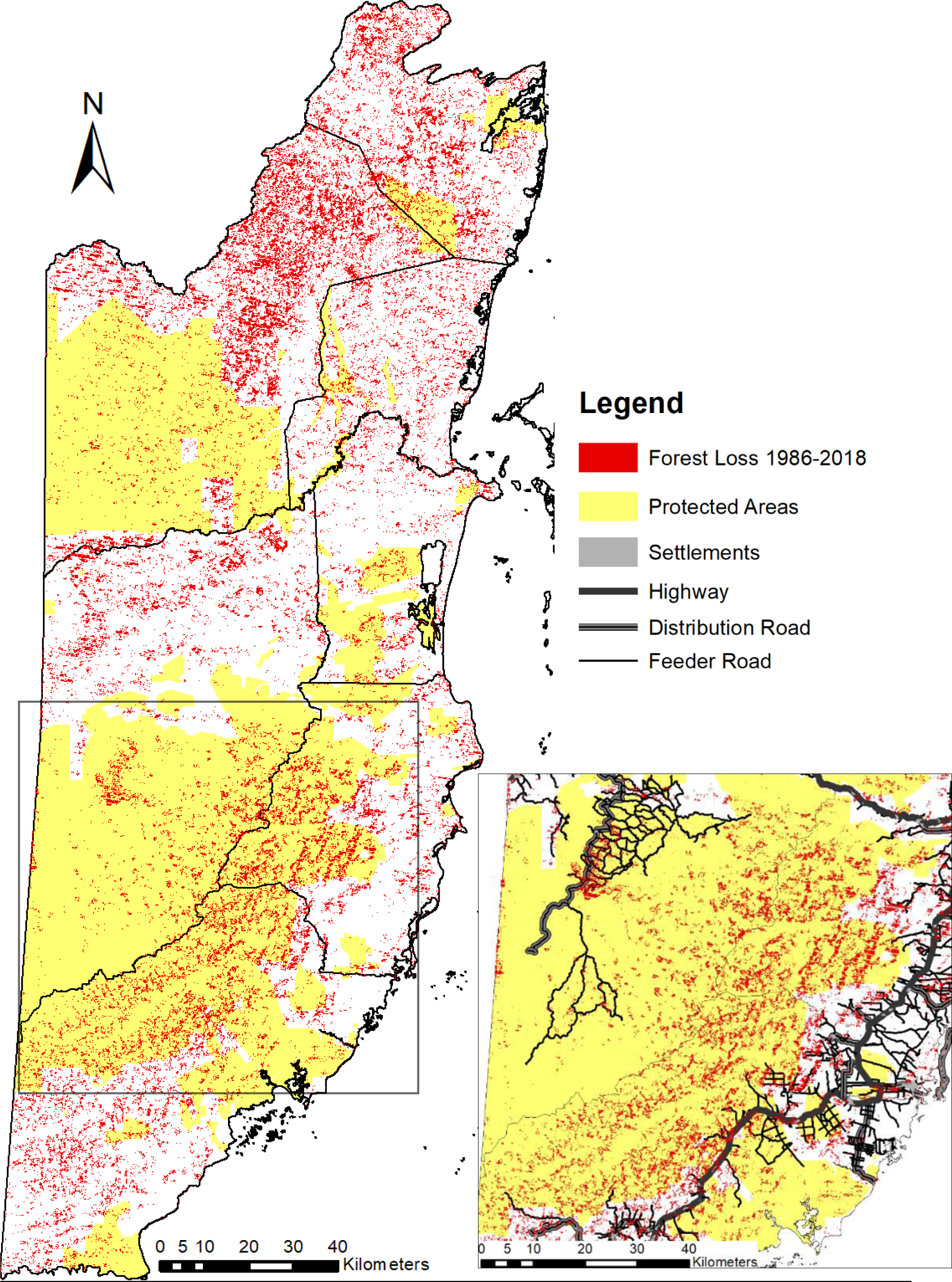
Areas of deforestation between 1986 and 2018, shown across protected and non-protected areas. Inset: a closer view of the largest protected area network including human infrastructure.

Of the three forest types experiencing the highest percentage of forest loss, only Ensino-Pixoy (Type 13a) occurs exclusively outside protected areas. Cohune Santa-Maria (Type 9a) and Sapote-Mahogany (Type 1) are found at least partially within protected areas and experience loss within them (Figure 7).

**Figure 7.**
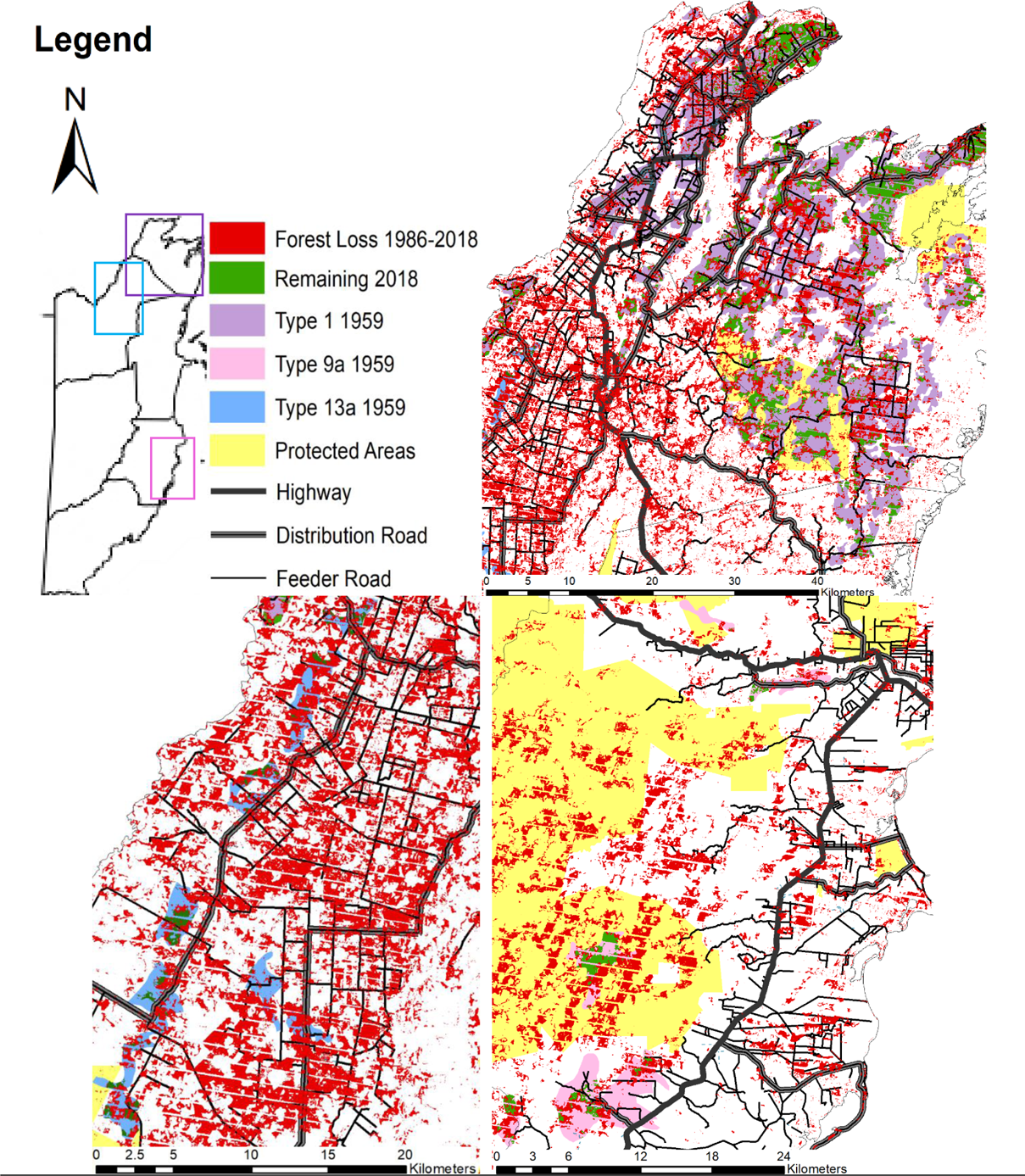
Habitat types with the highest percentage of deforestation. Top: Habitat Type 1, North Belize. Bottom left: Habitat Type 9a, Southeast Belize. Bottom right: Habitat Type 13a, Northwest Belize.

Mountain Pine Ridge is experiencing similar threats. Declared a protected forest in 1944, its designation changed to production forest in 1952, with intense timber exploitation commencing in 1959 (King *et al*., 1992). Before 1988 there was very little farming presence, with only small scale Milpa activities at the reserve’s northern boundary. The patch of intense deforestation visible in Figure 6 (inset) that coincides with road networks falls within the bounds of this park. Situated between the towns of San Antonio and San Ignacio, pressure for land development incurs on the forest. The reserve is known for its well-maintained and highly connective roads, which are maintained by the Forest Department and allow easy resource extraction, leading to higher levels of deforestation (Barber et al., 2014; Helmer et al., 2008; King *et al*., 1992). The Pinol soil type characterising this area is somewhat infertile and vulnerable to nutrient leaching and erosion, particularly on slopes of over 25% (King *et al*., 1992). This compounds the other threats to the area which include frequent forest fires, hurricane damage, and citrus farming, especially when ease of access is considered (Patterson, 2016; Voight, 2019).

In 1992 the National Resources Institute recommended that the reserve be “zoned into protection and production areas”, and “logging avoided in Pinol soil slopes” (King *et al.,* 1992). The high level of deforestation witnessed between 1986 and 2018, particularly in the areas with good road connectivity (Figure 8A), is evidence of these concerns not being heeded.

**Figure 8.**
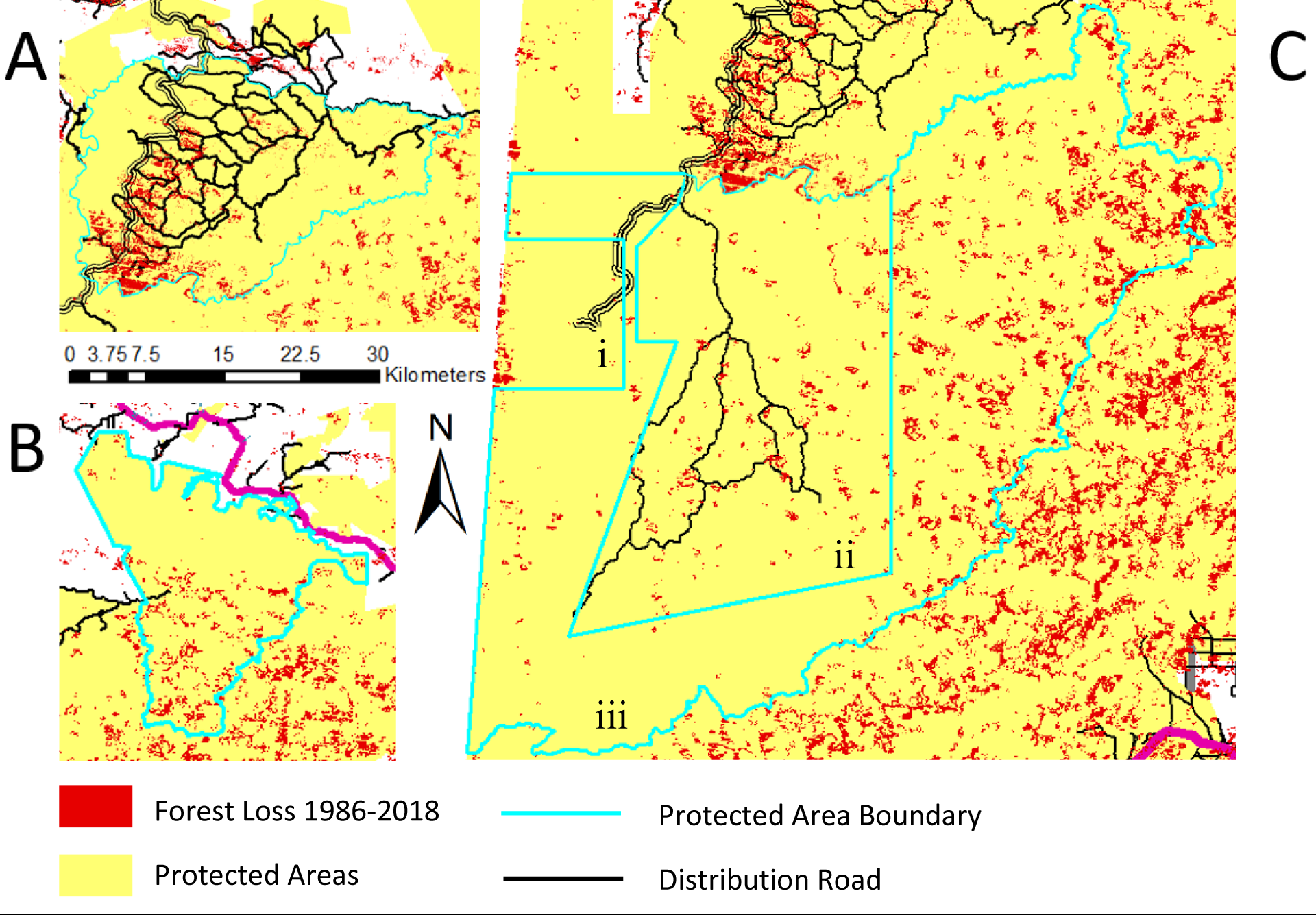
Protected areas experiencing high levels of deforestation. A) Mountain Pine Ridge Forest Reserve, B) Sibun Forest Reserve, C) i) Caracol Archaeological Reserve, ii) Chiquebul Forest Reserve, iii) Chiquebul National Park.

The Sibun reserve lies to the east of Mountain Pine Ridge. The northernmost lowland edge was heavily logged by 1992, partially aided by the creation of, and access granted by, the Hummingbird Highway (King *et al*., 1992). The rest of the reserve comprises of steep mountainous topography not conducive to production forestry. Therefore, the high incidence of forest loss detected towards the south (Figure 8B) is likely due to something other than timber extraction. Hurricane Iris of 2001 may be a cause, as the reserve lay within its path and was previously severely damaged by Hurricane Hattie in 1961 (Lewis, 2001; White *et al*., 2002).

An alternative explanation may be errors in land classification. One factor may be that as the imagery was taken from the peak of the dry season to minimise cloud cover, the vegetation ‘greenness’ and therefore its NDVI score may have fallen below 0.6 (Chicas *et al*., 2016), particularly as some species in these assemblages are semi-evergreen, seasonal forest (Wright *et al*., 1959). An even more likely explanation is relief and the resulting clouds. The topography in this area is high and steep, lying within the Richardson Peak, Mountain Pine and Xpiicila Hills land systems which provide clean water for most of the country (Mitchell *et al*., 2017). These ranges continue south into the Chiquebul Forest Reserve, where the arc of apparent deforestation matches the topography of the region (Figure 9).

**Figure 9.**
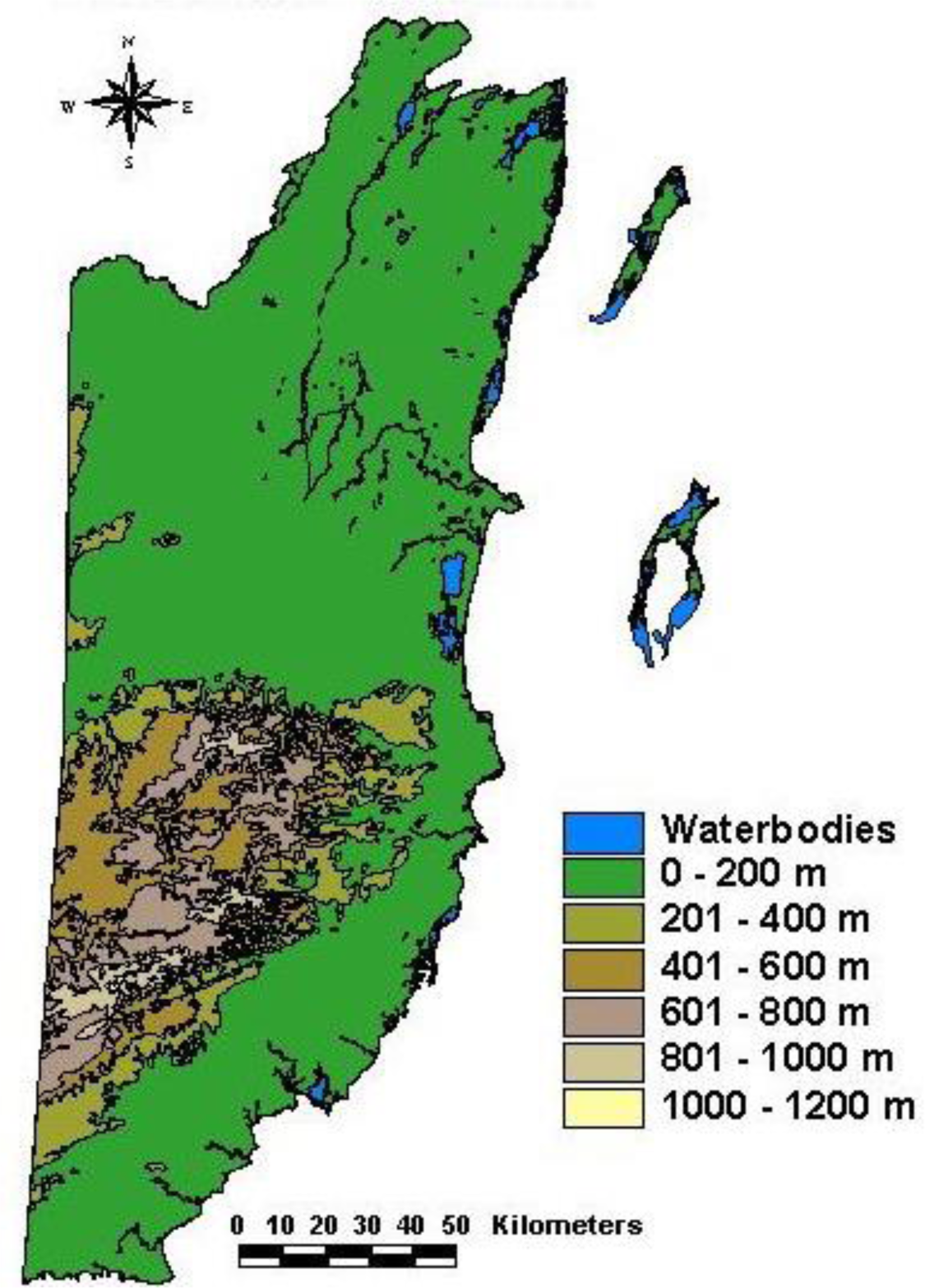
Elevations in Belize (Meerman, 2003)

While this may be attributable to seasonal changes in upland vegetation between 1986 and 2018, the pattern of cloud cover (Figure 3) indicates that clouds are the most likely explanation. The extremely high rainfall that southern Belize receives annually, around 4000mm, means that almost all days have at least partial cloud cover (King *et al*., 1986).

This study masked clouds themselves from the Landsat images, yet some of their shadows remained. Cloud shadow noise could not be entirely removed as the reflectance value in the red waveband was the same as some vegetation types. Removing them would have introduced bias against these species into the study. An alternative would have been to create a composite dataset from stacked Landsat images across many dates to maximise the chance of obtaining cloud-free pixels for the entire frame, as done by Meerman (2010). The issue with that method is that on different dates vegetation will be at different stages in its growth cycle, and so different ‘greenness’ will be observed despite there being no changes to forest stocking. To be accurate, change detection exercises should hold temporal, spatial, spectral, and radiometric resolutions constant across all timeframes (Jensen, 1996; Turner *et al*., 1989), and yet this is rarely possible, especially in the tropics (Kalacska, 2008). Whichever the chosen method, there will always be a compromise between precision, specificity and timeliness when using remotely sensed data (Harris and Ventura, 1995). This study’s methodology satisfies the requirement for temporal constancy as all three image frames for each year were from anniversary dates. Spatial resolution also remained constant at 30m for each image. The only spectral difference between Landsat 5 TM and Landsat 7 ETM+ is in the NIR band, where the TM uses 0.76-0.90 μm and ETM+ uses 0.77-0.90 μm. This difference is less than that in other studies with similar aims and methodologies (Emch *et al*., 2008), and Stow, Collins and McKinsey (1990) argue that change detection can still be highly accurate despite using satellites with different resolutions. In future studies, LiDAR and Radar could be used in place of Landsat imagery as they are less affected by cloud cover, but were not appropriate as the data source for s study as imagery only dates back to 1991 (Emch *et al*., 2008). However, Radar and LiDAR can be used in addition to the deforestation data generated by this study to explain recent deforestation in specific areas.

Even in regions that are under full protection, where agriculture and forest clearance are forbidden, lack of financial resources and on-the-ground management allow poaching to continue. The term ‘paper parks’ describes areas that have legal designations but lack the necessary resources to enforce their protection (Blackman *et al*., 2015).

Some areas of detected deforestation within the Chiquebul cannot be attributed to cloud shadow errors, despite its protected status. Together, the Caracol Archaeological Reserve, Chiquebul Forest Reserve, and Chiquebul National Park (Figure 9) form part of the largest contiguous expanse of neotropical forest north of the Amazon (Weishampel *et al*., 2012). Pockets of deforestation are visible along the Guatemala-Belize border, especially within the Caracol Archaeological Reserve. Despite only a few sustainable logging practices permitted since 1994, the majority of deforestation in this area has been found to have occurred since 2000 (Weishampel *et al*., 2012). This could not have been caused by hurricanes, as the last significant storm to affect the area was Hurricane Greta in 1978. The pattern of deforestation exactly follows the southern boundary of the reserve, indicating it is caused intentionally by humans, and that management of the park is not effective (Figure 8Ci).

Among the Guatemalan poor persists a belief that Belize is their rightful territory (Young, 2008). This leads to poaching, harvesting of forest products and looting of historical artefacts that although illegal in Belize, the people of Guatemala consider to be their right. Deforestation from these activities has been observed to occur as far as 2km east of the border due to a lack of conservation posts around the Caracol Reserve (Weishampel *et al*., 2012). Belizean authorities are attempting to remedy this ‘paper park’ situation by building new ranger outposts, yet threats from Guatemalan farmers complicate the issue (Chicas et al., 2016). In 2014, murder and armed conflicts occurred at the Mayan site between Guatemalan subsistence farmers and members of the Belize Defence Force over plans for increased conservation efforts at Valentin Camp, Cayo, a hotspot for Guatemalan incursion and poaching (Ramos, 2014). This area is used for the illegal agriculture including farming of maize, pumpkin and beans (Figure 10), as well as being an entry point for illegal loggers prospecting further into the Chiquebul National Park and Forest Reserve (Ramos, 2014). In addition to loss of flora, poaching of fauna is a concern at Caracol. The reserve is home to kinkajou, *Potos flavus*, and black howler monkeys, *Alouatta pigra*, whose populations are nationally at risk due to habitat fragmentation and conversion to agricultural land (Estrada *et al*., 2006).

**Figure 10.**
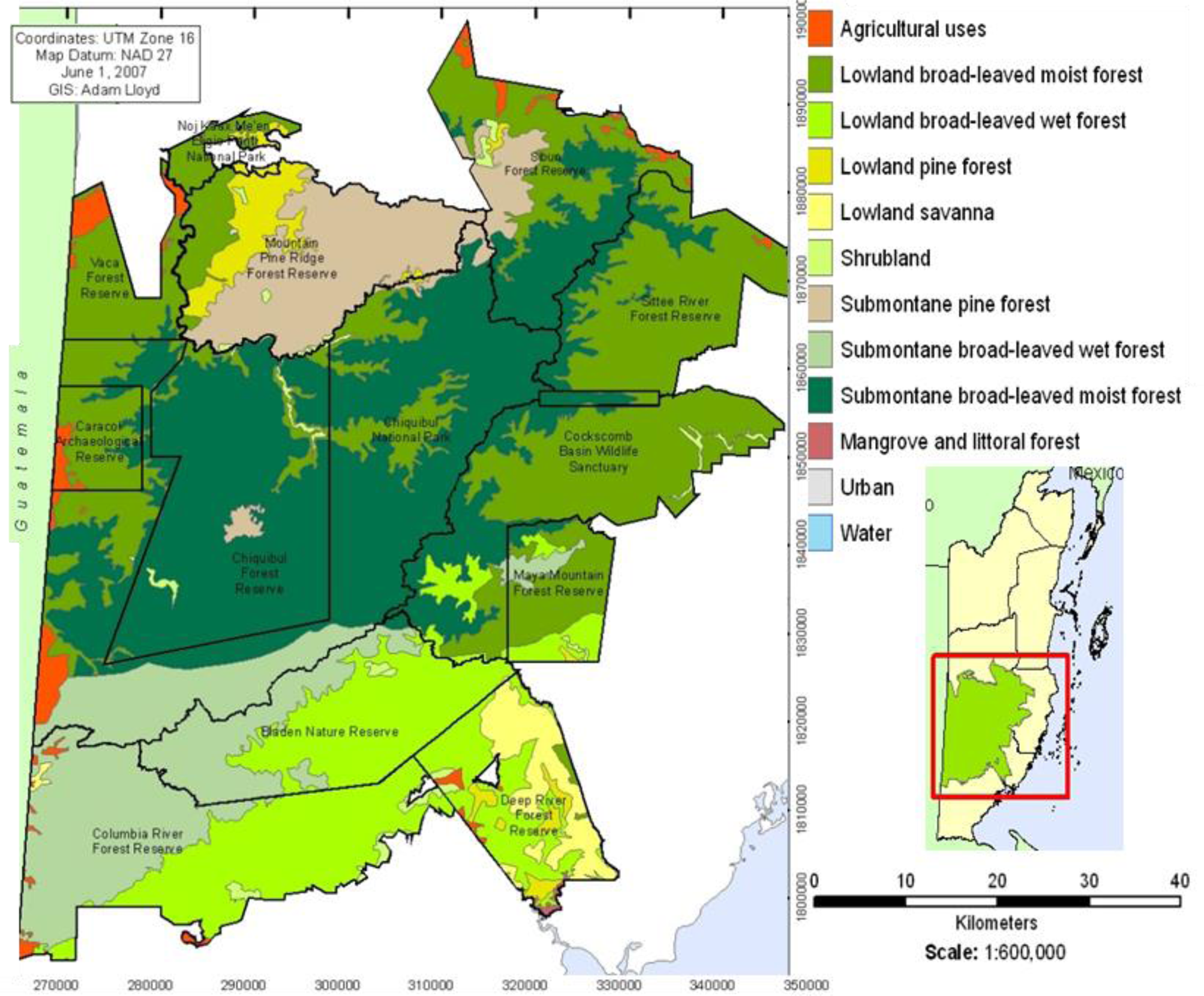
Protected areas of the Maya forest, Central Belize, showing areas natural vegetation, and historical and modern agricultural use (Lloyd, 2007).

The Caracol ruins were discovered in 1937 by a logger prospecting for Mahogany (Chase and Chase, 1987), which was fairly abundant in this region in the 20^th^ Century (Wright *et al*., 1959). The archaeological site varies in tree species composition from the majority of the Chiquebul area, comprising of Semi-evergreen Chiquebul-Ramon (Type 3), Ramon-Chiquebul (Type 3a) and Ironwood-Chiquebul (Type 3b) habitats (Figure 8Ci). Ramon, commonly known as Breadnut, is strongly associated with Mayan sites and agriculture (Puleston, 1978), as it was a staple food source grown in Maya forest gardens (Ford, 2008) (Figure 9). These three habitat types are among those experiencing the smallest percentage loss from 1986 to 2018 (Figure 4). In only one location does any of these Mayan forests abut the Guatemalan border; a medium-sized fragment of Chiquebul-Ramon. In all other instances, the patches are at least 2km from the border, which is the limit of Guatemalan-driven deforestation detected in previous studies (Weishampel *et al*., 2012). The proximity of these forest types to tourist activity and academic activity may be protecting them from the levels of deforestation experienced by surrounding areas. The unsuitability of the single-track Caracol road for agricultural or forestry traffic also reduces ease of poaching behaviours in this area (Wyman, 2008). Plans to use OPEC funding to upgrade this 26-mile track to a two-lane, all-weather track costing $180 million (Sanchez, 2019; Taegar-Panton, 2019) may have a detrimental effect by increasing ease of access for poachers as well as tourists and legitimate farmers. This possibility is in addition to concerns over the size of the loan and associated interest rate when the money could be better spent improving rural quality of life and reducing poverty, both of which often lead to poaching activities (Amandala, 2019).

Chiquebul-Cherry (Type 2d) forest, which surrounds many of these Ramon-rich fragments, remains the most abundant of all vegetation types for all timeframes due to its wide distribution, its area reduced by only 17% between 1987 and 2018. It is the primary constituent of the Chiquebul Forest Reserve and National Park, and occurs solely within these areas. This common habitat is found in large, uninterrupted swathes extending deep into the central areas of the reserve, where distance from roads and unprotected borders acts as a natural defence against deforestation. This shows the importance of large, contiguous protected areas where the influence of edge effects is diminished and natural barriers aid human efforts in protecting economically and ecologically valuable species. The most significant fragmentation Chiquebul-Cherry experiences is along the Guatemalan border.

Chiquebul-Cherry and the eight other Lime-Loving Broadleaf forests that comprise the majority of the Chiquebul Forest Reserve host the high-value timber species Mahogany and Cedar, supplemented by Santa Maria, Fiddlewood, Nargusta, Mylady, Billywebb and Ironwood (Wright et al., 1959). Of these, Mahogany (VU) Cedar (VU) and Fiddlewood (EN) are classified by the IUCN Redlist (Mark and Rivers, 2017). The reserve has been exploited for these species since 1925, and only received reserve designation in 1956 (King *et al*., 1992). Wright *et al*. (1959) noted the occurrence of Mahogany and Cedar in each Chiquebul habitat type down to the average number per acre. This was generally low, around one to two or three acres, except in rare pockets of higher abundance. By this point in time, forest stocks had been depleted, and further logging and the occurrence of hurricane Hattie in 1961 made the stocking extremely poor by 1971 (King *et al.,* 1992). In 1989, estimates of regeneration declared that the number of young stands likely to establish and succeed was low (ODA, 1989). Logging was once again permitted in the north-west corner of the reserve, despite recommendations to allow recovery (King *et al*., 1992).

No Chiquebul forest type was found to experience over 50% loss from 1986 to 2018, indicating that despite threats from the border, management of this reserve is somewhat effective. Of the Chiquebul forest types, Santa Maria-dominated habitats experienced the highest loss from, with Type 4 at 47%, and 4b at 41% loss. This may be due to Santa-Maria’s uses as both a medicine and a high-value timber species, primarily for production of furniture, decking, flooring and veneer (WWF, 2013). Santa Maria is classified by the IUCN as Least Concern (BGCI, 2018), and is suggested by WWF as an alternative species to White Ash (CR), Sapele (VU), Mahogany (VU), African Mahogany (VU) and African Walnut (LC) (WWF, 2013). Santa-Maria dominated forests are also threatened outside of the Chiquebul. Further east, habitat type 9a is being lost from Mango Creek Forest Reserve. (Figure 7).

It is possible that Santa-Maria is not the target species when deforestation occurs in these areas. Forest types 4 and 4b also once boasted a density of mahogany and cedar of approximately one to the acre- the highest density of any Chiquebul type- and local pockets might have hosted up to three per acre (Wright *et al*., 1959). In addition to Mahogany, Santa-Maria habitats also host *Heliconia* species; a popular ornamental plant which may provide daytime habitat for bats (Eklöf and Rydell, 2018), last classified as Vulnerable in 2004 (Ulloa Ulloa and Pitman, 2004).

Only a small number of Belizean taxa have received a conservation status from the IUCN, with the abundance of some high-value species unknown. Though no record is held by the IUCN, Belizean Rosewood *Dalbergia stevensonii* is a CITES species still suffering poaching and illegal trade, particularly export to Asian markets (Wainwright and Zempel, 2018). This demand for Rosewood along with a population increase and weak enforcement has caused a rise in deforestation in the protected areas of Toledo (Chicas *et al.,* 2016), which abut the Chiquebul in the north of the district. In 2012 the issue became so extreme that the Government declared a moratorium on remaining legal exports of Rosewood lumber (Ya’axche, 2013).

New threats to the forests of Belize have altered the reasons for the designation of protected areas from controlling unsustainable logging when the industry was at its peak to balancing the effects of increasing human population and its demand for land and resources (Young, 2008). Yet, it appears that the emergence of these new threats has not replaced the old, only compounded them. This study found level of deforestation in protected areas to be only slightly less than the national average. This follows the prediction made by Amor and Christensen (2008) that there is no clear difference in the likelihood of deforestation occurring inside or outside of Belize’s predicted areas. Across all protected areas, 83,361 ha was lost, representing 35% of all forest loss. Once again this varies from Cherrington and Ek’s (2010) findings, in which only 15.2% of the deforestation from 1980-2010 was seen to occur in protected areas. The disparity in these calculations may be due once more to differences in determining secondary forest growth, or a rise in deforestation rate, particularly focussed in specific areas such as Caracol or the Northern plains (Figure 7). The other consideration is the areas of deforestation measured along the mountains, which artificially inflate the figure. It is likely a combination of these factors, varying between reserves with different levels of protection and valuable species. Further study including ground-truthing of remote sensing data will be required to determine the exact trend. This is vital as the management of protected areas in Belize is restricted by limited resources. If current management strategies are not effective, then the most vulnerable habitats must be identified and prioritised in terms of law enforcement and community support. Where this applies to the Guatemala border, bilateral partnerships between the governments of both countries, with input from NGOs and local communities, are necessary to ensure that management strategies are effective and do not lead to further violence.

To attain a more detailed understanding of *what* is being lost, ground studies are again necessary to determine if the Wright *et al*. 1959 forest types are still accurate in terms of distribution and species composition, or if a new vegetation survey is required to understand the abundance of key species. As part of this, a thorough review of species indigenous to Mesoamerica is needed to update and complete IUCN records of these taxa, as currently the database is data deficient.

The *where* and the *why* of deforestation in Belize have been answered more thoroughly. This study is the first to provide both a quantitative country-wide assessment of deforestation by vegetation type and its leading causes. This sets a baseline of forest cover in 2018 which can be used by future studies to assess ongoing effectiveness of protected areas and advise policy, as well as aiding other applications such as calculation of carbon stocks or ecosystem service integrity.

## Appendix 1

IDEFO diagrams depicting detailed methodology for image processing. IDEF0 is a function modelling technique commonly used for software engineering and business processes. In this case it is useful for modelling the decisions, actions, and activities involved in this non-automated process (Lightsey, 2001).

**Figure 1:**
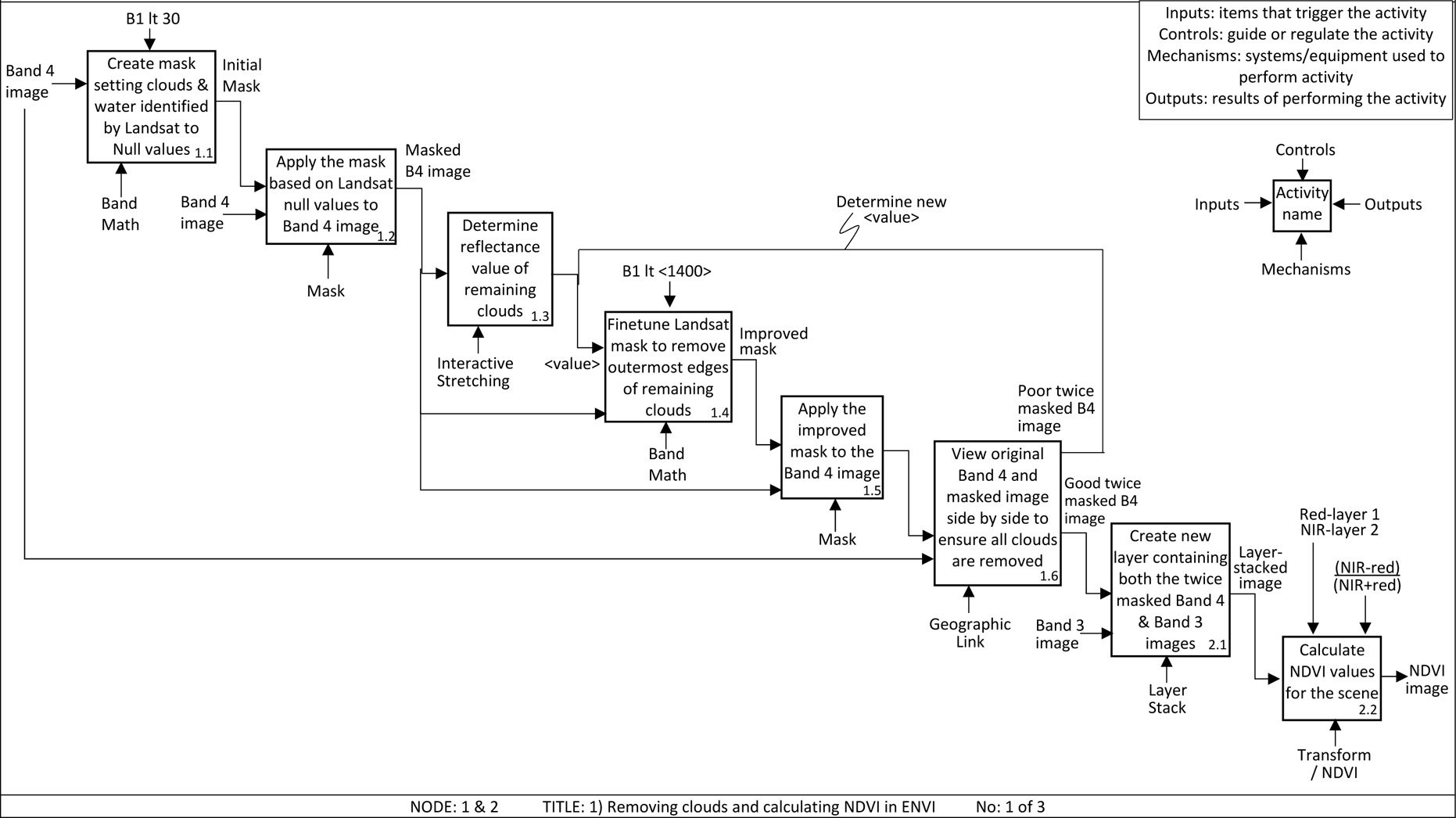
IDEF0 diagram detailing the stages of image pre-processing Landsat data in ENVI.

**Figure 2:**
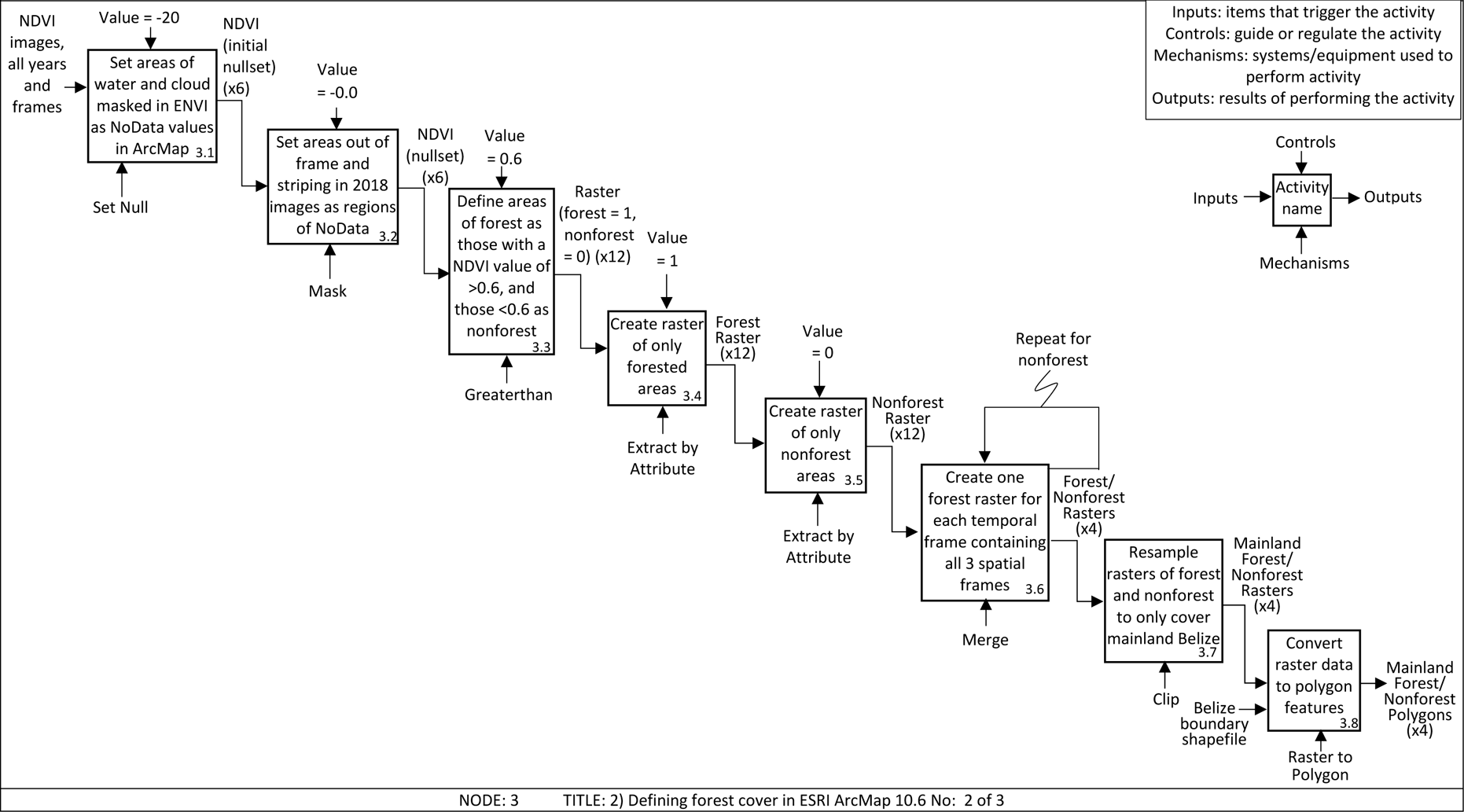
IDEF0 diagram detailing the stages of classifying forest/nonforest areas from Landsat NDVI data in ArcMap 10.6.

**Figure 3:**
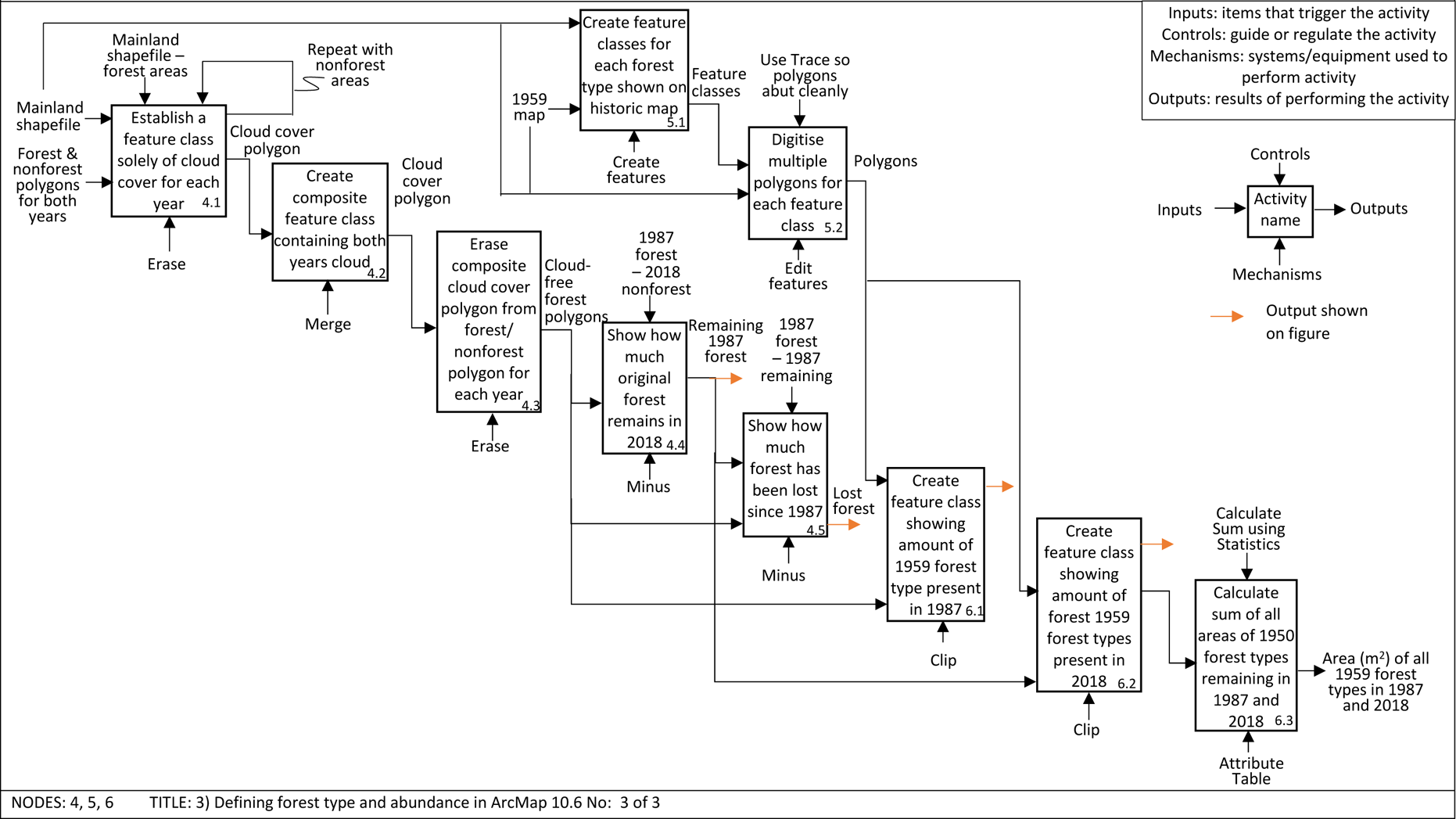
IDEF0 diagram detailing the process of defining forest type from historic maps and calculating their abundance in ArcMap 10.6.

## Appendix 2

**Table 1.**
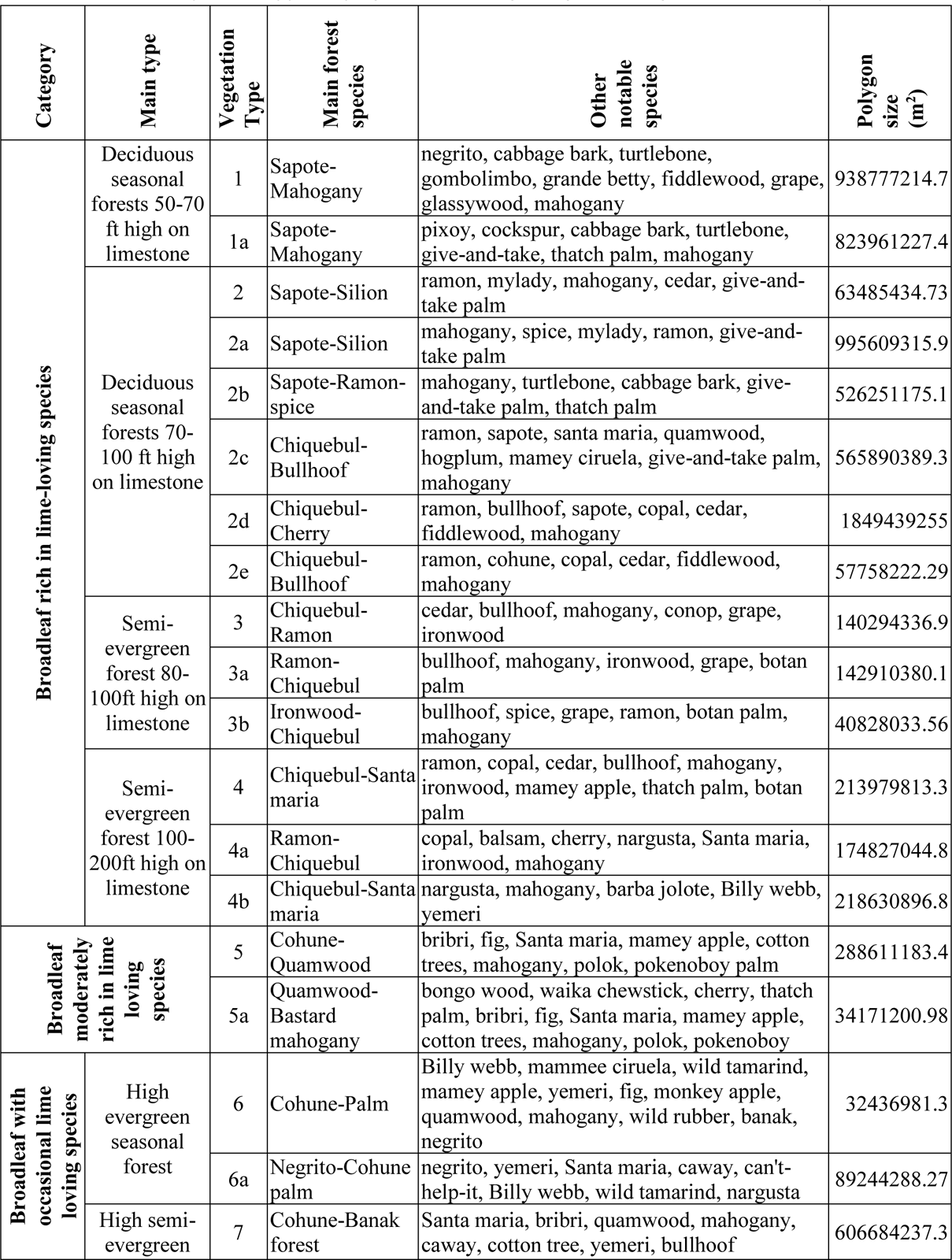
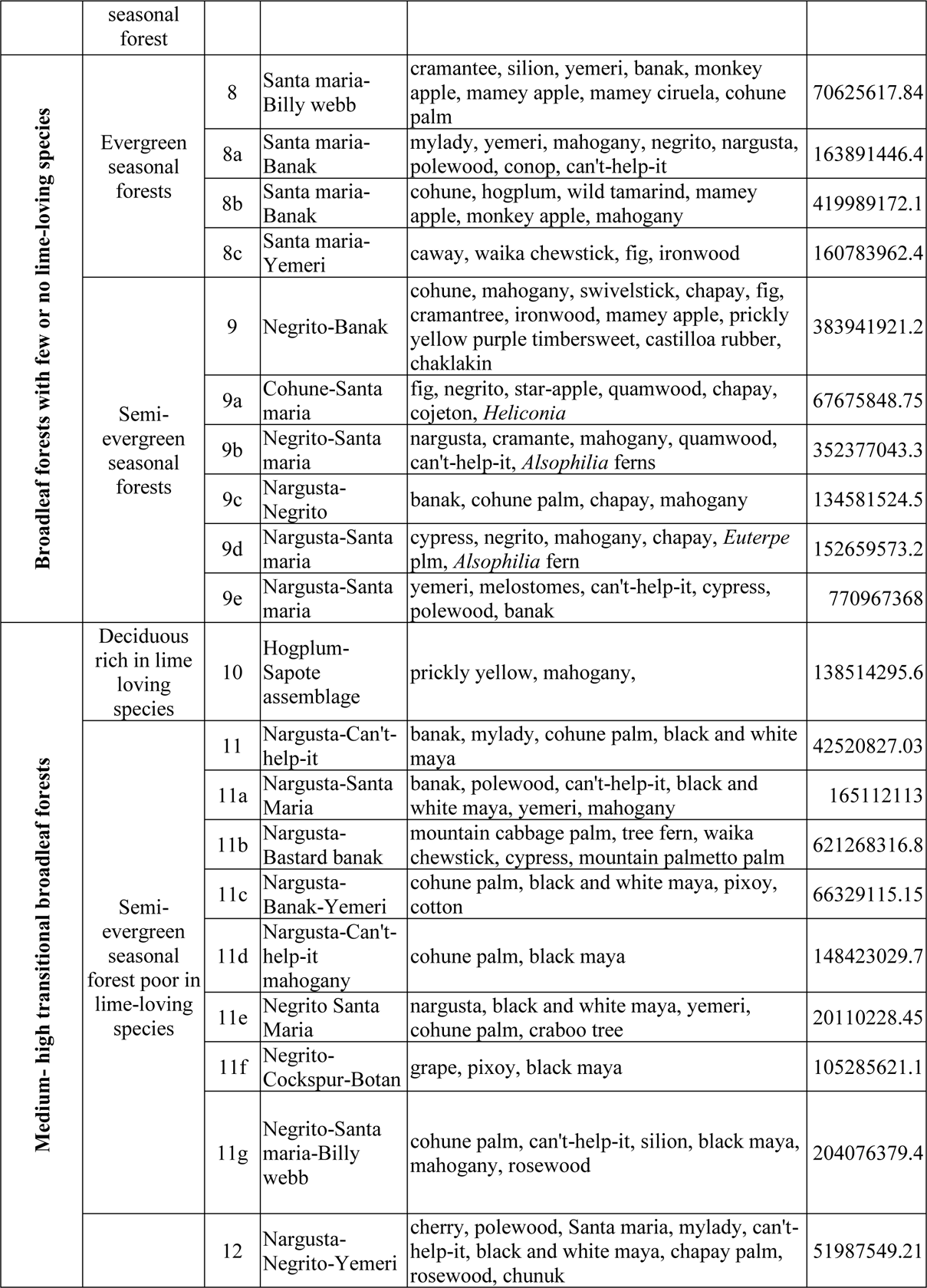
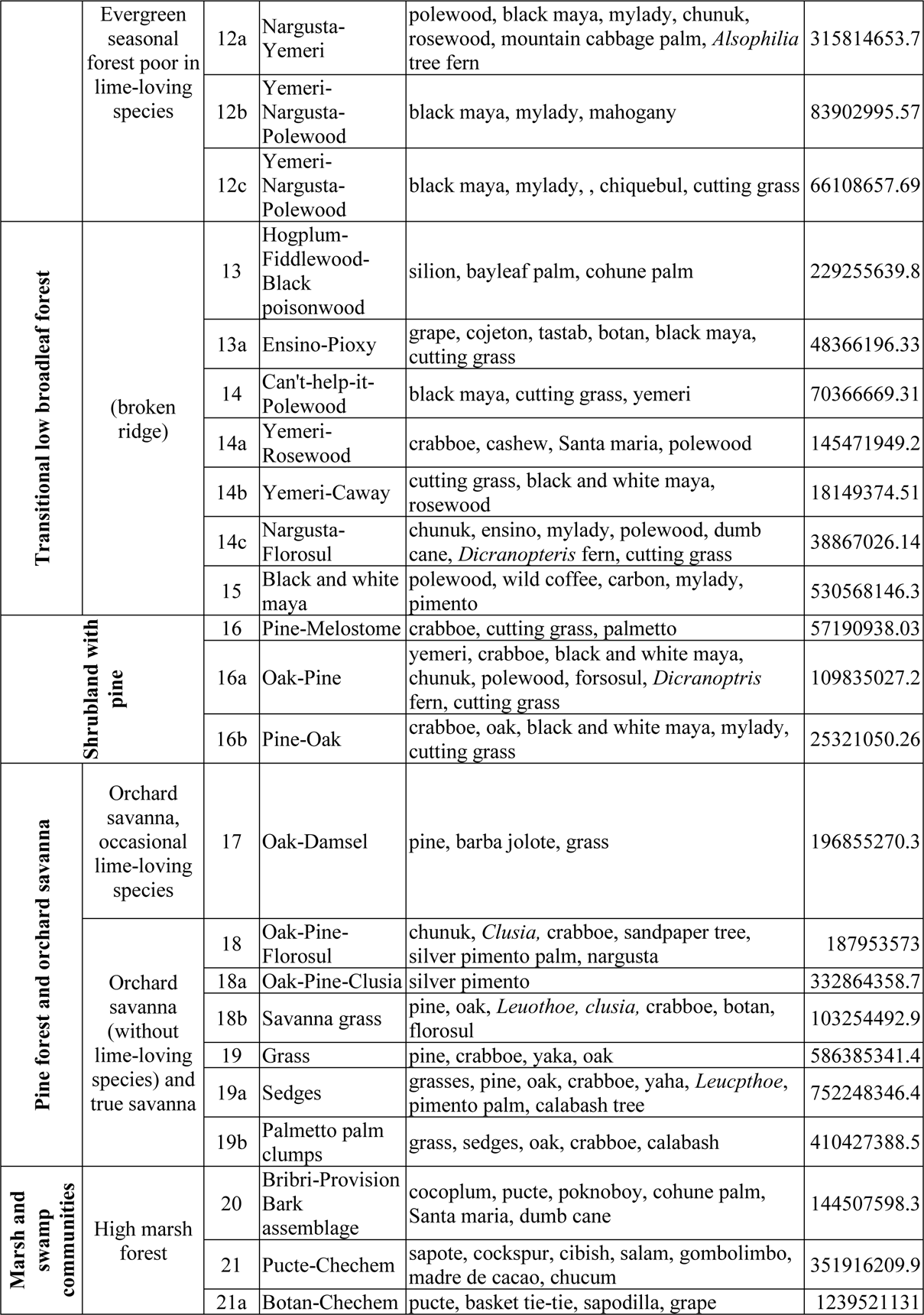
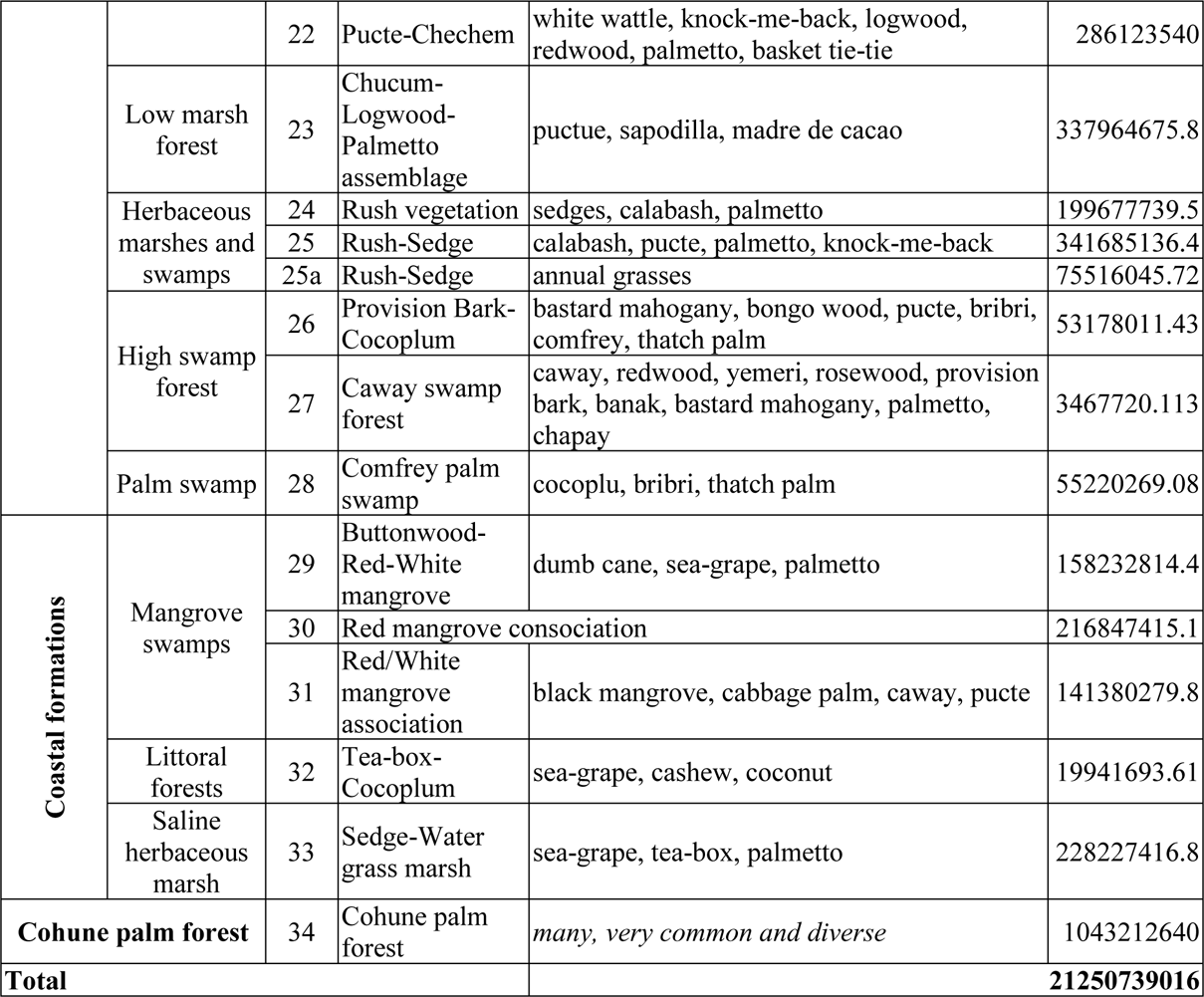
Size and composition of forest fragment according to digitised Wright et al. 1959 maps

